# SARS-CoV-2 detection with CRISPR diagnostics

**DOI:** 10.1101/2020.04.10.023358

**Authors:** Lu Guo, Xuehan Sun, Xinge Wang, Chen Liang, Haiping Jiang, Qingqin Gao, Moyu Dai, Bin Qu, Sen Fang, Yihuan Mao, Yangcan Chen, Guihai Feng, Qi Gu, Liu Wang, Ruiqi Rachel Wang, Qi Zhou, Wei Li

## Abstract

The novel coronavirus (CoV) disease termed COVID-19 (Coronavirus Disease-19) caused by SARS-CoV-2 (Severe Acute Respiratory Syndrome Coronavirus-2) is causing a massive pandemic worldwide, threatening public health systems across the globe. During this ongoing COVID-19 outbreak, nucleic acid detection has played an important role in early diagnosis. Here we report a SARS-CoV-2 detection protocol using a CRISPR-based CRISPR diagnostic platform - CDetection (Cas12b-mediated DNA detection). By combining sample treatment protocols and nucleic acid amplification methods with CDetection, we have established an integrated viral nucleic acid detection platform - CASdetec (CRISPR-assisted detection). The detection limit of CASdetec for SARS-CoV-2 pseudovirus is 1 × 10^4^ copies/mL, with no cross reactivity observed. Our assay design and optimization process can provide guidance for future CRISPR-based nucleic acid detection assay development and optimization.

## Results

The novel coronavirus (CoV) disease termed COVID-19 (Coronavirus Disease-19) caused by SARS-CoV-2 (Severe Acute Respiratory Syndrome Coronavirus-2) [1] is causing a massive pandemic worldwide, threatening public health systems across the globe. During this ongoing COVID-19 outbreak, nucleic acid detection has played an important role in early diagnosis [2]. To date, 4 protocols based on CRISPR for detecting SARS-CoV-2 have been published [3–6]. Using lateral flow protocols, RNA samples harboring more than 1 × 10^4^ - 1 × 10^5^ copies/mL (SHERLOCK) or 1 × 10^4^ copies/mL (DETECTR) can be detected within 1 hour. In addition to these reported efforts, we have also established a SARS-CoV-2 detection protocol based on our previously reported platform – CDetection (Cas12b-mediated DNA detection) [7]. By combining sample treatment protocols and nucleic acid amplification methods with CDetection, we have established an integrated viral nucleic acid detection platform – CASdetec (CRISPR-assisted detection). The detection limit of CASdetec for SARS-CoV-2 pseudovirus is 1 × 10^4^ copies/mL, with no cross reactivity observed. Here we present our assay design and optimization process, which could provide guidance for future CRISPR-based nucleic acid detection assay development and optimization.

To optimize the output of fluorescence signal, we designed and synthesized poly-T fluorescence-quenchers of varying nucleotide lengths, namely 4 nt, 5 nt, 7 nt, 12 nt, 17 nt, 22 nt and 27 nt. Of all the lengths tried, the 7 nt poly-T reporter provided the highest signals in the shortest amount of time (Supplementary information, Fig. S1a, b). Based on this observation, we applied the 7 nt poly-T reporter in later experiments.

According to the published SARS-CoV-2 whole genome sequence [8], we designed 7 sgRNAs around the RdRp locus, as recommended by the World Health Organization (WHO) [2] (Supplementary information, Fig. S2a). Due to its high similarity to SARS-CoV, we ran initial experiments on both SARS-CoV-2 and SARS-CoV plasmids. According to fluorescence kinetics studies, sgRNA-3 stood out in not only being able to distinguish between the 2 similar coronaviruses but also being able to produce the most distinct fluorescence signal (Fig. 1a and Supplementary information, Fig. S2b-g).

**Figure 1.**
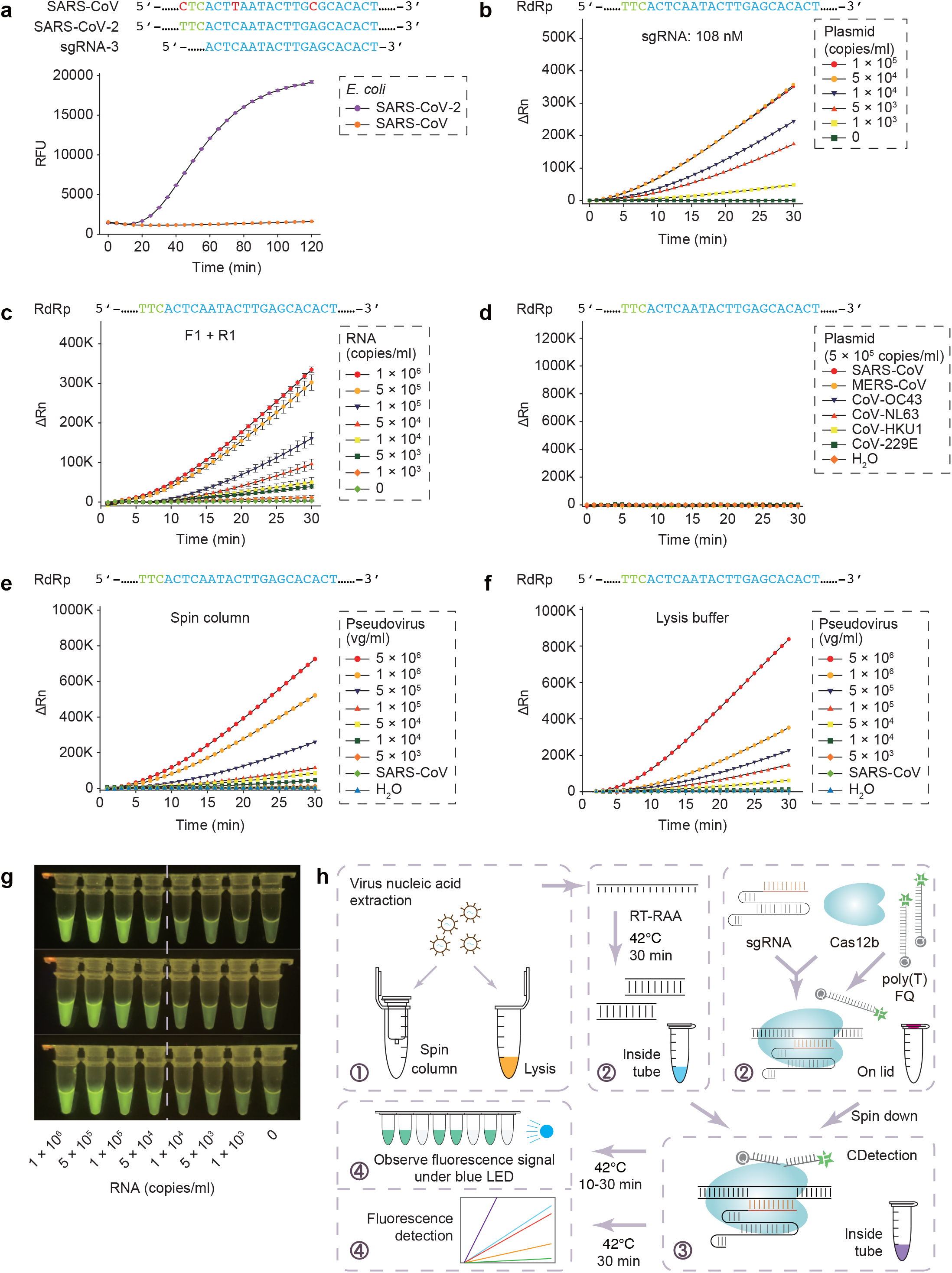
CASdetec used for SARS-CoV-2 detection. **(A)** Fluorescence kinetics of sgRNA-3 for RdRp detection. *E. coli* cells bearing Blunt-SARS-CoV-RdRp or Blunt-SARS-CoV-2-RdRp were pre-incubated at 95□ for 10 min and used as templates for RAA and CDetection. PAM sequences are colored in green, protospacers are colored in blue, base pair mismatches are colored in red. Error bars indicate standard errors of the mean (s.e.m.)., *n* = 3. RFU, relative fluorescence units. **(B)** Fluorescence kinetics of RdRp detection using 108 nM sgRNA-3. Plasmid bearing SARS-CoV-2-RdRp was serially diluted as shown in the legend. *n* = 2. ΔRn, ΔFluorescence, which refers to the Rn value of an experimental reaction minus the Rn value of the baseline signal generated by ABI 7500. **(C)** Fluorescence kinetics of F1- and R1-based RdRp detection. SARS-CoV-2-RdRp RNA was serially diluted as shown in the legend. Error bars indicate (s.e.m.), *n* = 3. **(D)** Evaluation of cross reactivity. Plamids containing target RdRp region from 6 human epidemic coronaviruses are serially diluted as the shown in the legend. *n* = 2. **(E)** Detection of SARS-CoV-2 pseudovirus. Virus genome is extracted using the virus RNA extraction kit (spin column). SARS-CoV were diluted to 5 × 10^5^ copies/mL. *n* = 2. **(F)** Detection of SARS-CoV-2 pseudovirus. Virus was treated by direct lysis. SARS-CoV was diluted to 5 × 10^5^ copies/mL. *n* = 2. **(G)** CASdetec results can be directly observed under blue LED. 3 replicates of product from Fig. 1b are imaged upon blue LED illumination. **(H)** Schematics showing the workflow of CASdetec. Virus genome is extracted by kit or direct lysis. Target sequences are pre-amplified by isothermal amplification, followed is CDetection. Fluorescence signals are obtained either from fluorescence reader or direct observation under blue light.

As previously demonstrated, CRISPR is unable to detect any target DNA when there is less than 1-10 nM of amplification product within the reaction mix [7]. Hence, increasing the molecular collisions between CRISPR and target would be essential to improve sensitivity. We found out that increasing the sgRNA concentration by 3 folds not only enhances the fluorescence signal and the signal-to-background ratio, but also increases the rate of reaction (Fig. 1b and Supplementary information, Fig. S3a-b).

Given that the average viral load in the plasma of SARS patients ranged from less than 1 to about 1000 copies per microliter [9], or 1 × 10^3^ – 1 × 10^6^ copies/mL, nucleic acid amplification techniques are needed to produce sufficient DNA for CRISPR-based DNA detection methods. Recombinase-aided amplification (RAA) can amplify substrates 10^10^ times at most (from aM to 10 nM) within 10-30 minutes at constant temperature between 37°C to 42°C, complementing the needs of CRISPR-based detection. Thus, we designed and screened RAA primers that matched our previously optimized sgRNA-3 (Supplementary information, Fig. S4a). Based on our screens, we found that by using the best primer pairs together with sgRNA-3, we can detect SARS-CoV-2 RNA in samples containing 5 × 10^3^ copies/mL (Fig. 1c and Supplementary information, Fig. S4b-c).

In addition, we aligned the selected primers and sgRNAs to the existing typical coronavirus sequences to evaluate their specificity. We analyzed all SARS-CoV-2 sequences that have been uploaded to GISAID up till March 26^th^ 2020. Out of 1792 sequences on GISAID, 1673 of them contained sequences matching our chosen primers and sgRNA. Only 3 of them have 1 mismatch to the forward primer and only 2 of them have 1 mismatch to the reverse primer (Supplementary information, Fig. S5), suggesting that our selected sgRNA and primers can be used for nearly all of the reported SARS-CoV-2 genomes. Meanwhile, we aligned 12 typical human coronaviruses, and found that none of the whole set of primers and sgRNA showed high similarity (Supplementary information, Fig. S6a-c).

The experiments above were conducted by executing RT-RAA nucleic acid amplification and CDetection separately. However, it would be best to conduct both reactions within a single tube for convenience and, more importantly, to prevent aerosol contamination which happens when the reaction mixture has to be exposed to the environment midway through the protocol. Hence, we tried to execute both the RT-RAA and CDetection concurrently within a single tube. However, the combination resulted in a drastic decrease in sensitivity (Fig. 1c and Supplementary information, Fig. S7). Therefore, in order to keep both the RT-RAA and CRISPR reactions within a single tube, we executed the RT-RAA reaction within the tube while keeping the CDetection reagents within the lid of the tube for 30 minutes, following which, the CDetection reagents were spun down into the tube for nucleic acid detection, and the resultant reaction mixture was imaged for fluorescence.

To validate the specificity of our method for SARS-CoV-2 nucleic acid detection, we tested our protocols against 6 coronaviruses known to cause respiratory diseases (SARS-CoV, MERS-CoV, CoV-HKU1, CoV-229E, CoV-OC43 and CoV-NL63). Consistent with alignment analysis (Supplementary information, Fig. S6a-c), no cross-reactivity with other endemic human coronavirus were detected (Fig. 1d). Our results suggested our set of sgRNA and primers showed high sensitivity and specificity.

However, viral genomes are packaged inside capsid protein and need to be released. Thus, to investigate the virus handling processes, we produced pseudoviruses by packaging the target sequences of SARS-CoV-2, SARS-CoV, MERS-CoV into actual lentivirus particles (Supplementary information, Fig. S8). These pseudoviruses were diluted serially and treated with either virus genome extraction kits (spin column) or lysis buffer, respectively. Our results demonstrated that the spin column treatment gave a lower detection limit of 1 × 10^4^ copies/mL (Fig. 1e). On the other hand, the lysis buffer offered higher usability, but raised the detection limit to 5 × 10^4^ copies/mL (Fig. 1f). Due to the difference in detection limit between spin column and lysis buffer, we suggest using the spin columns in hospitals, and using the lysis buffer for point-of-care testing (POCT).

To make CASdetec more amenable for POCT, we have also constructed a portable dark box containing a blue LED and demonstrated that the positive fluorescence signal generated from the protocol can be visualized upon illumination by a blue LED (Fig. 1g). In conclusion, we have established a CRISPR-assisted detection (CASdetec) platform which consists of procedures including virus handling, nucleic acid amplification and CRISPR-based detection (Fig. 1h). CASdetec can detect pseudovirus samples with more than 1 × 10^4^ copies/mL, with no cross-reactivity with other endemic human coronaviruses. In addition, we optimized the workflow to run both reactions within one single tube without lid opening. This will thus prevent aerosol contamination and reduce the false positive rate.

## Acknowledgments

We thank Prof. Ng Shyh◻Chang from Institute of Zoology, CAS for his critical support with this study. We thank Hanxing Zhang from Institute of Microbiology, CAS for her kind help on equipment. This work was supported by the National Key Research and Development Program (2020YFA070009 to R.R.W), the Key Research Projects of the Frontier Science of the Chinese Academy of Sciences (QYZDY-SSW-SMC002 to Q.Z. and QYZDB-SSW-SMC022 to W.L.), the Strategic Priority Research Program of the Chinese Academy of Sciences (XDA16030400 to W.L.).

## Author contributions

L.G., R.R.W., Q.Z., and W.L. conceived and designed the experiments. L.G., X.S., X.W., C.L., H.J., Q.G., M.D., B.Q., S.F., Y.M. and Y.C. participated in multiple experiments; L.G., X.W., X.S., C.L., and G.F. analyzed the data. L.G. wrote the manuscript. W.L., R.R.W., Q.Z., and L.W. provided the final approval of the manuscript.

## Notes

### Competing Interest Statement

The authors have declared no competing interest.

