## Supplementary information, materials and methods for "SARS-CoV-2 detection with CRISPR diagnostics"

***Protein purification***

AaCas12b proteins were purified by GenScript. Briefly, BPK2014-AaCas12b-$His_{10}$ was cloned into *E. coli* strain BL21 (λDE3) and expression was induced with IPTG at 15 °C for 16 h. Cell pellets were resuspended with lysis buffer followed by sonication. Target protein was obtained by two-step purification using Ni column and Superdex 200 column. Purified AaCas12b proteins were dialyzed, concentrated and quantified using BCA Protein Assay Kit (Thermo Fisher).

***Nucleic acid preparation***

DNA oligos were commercially purchased (GenScript). Double-stranded DNA activators were obtained by PCR reaction and purified using Oligo Clean & Concentrator Kit (ZYMO Research). In order to avoid false positive results caused by target strand (TS) ssDNA, we used non-target strand (NTS) ssDNA as PCR template. PCR primers and ssDNA templates were listed in Supplementary information, Table1.

Guide RNAs were transcribed *in vitro* using HiScribe™ T7 High Yield RNA Synthesis Kit (NEB) and purified using MicroElute RNA Clean Up Kit (Omega). AaCas12b sgRNA (AasgRNA) templates for *in vitro* RNA transcription were PCR amplified using primers bearing a T7 promoter (Supplementary information, Table1).

Target sequences were assembled by high-fidelity PCR as previously reported [1]. Briefly, DNA oligos were commercially purchased (GenScript) (Supplementary information, Table1). PCR products were ligated to pEASY-Blunt (TransGen) and sequenced. Plasmids were extracted from *E. coli* clones bearing the right sequences.

SARS-CoV-2-RdRp RNA was transcribed *in vitro* using HiScribe™ T7 High Yield RNA Synthesis Kit (NEB) and purified using MicroElute RNA Clean Up Kit (Omega). SARS-CoV-2-RdRp RNA templates for *in vitro* RNA transcription were PCR amplified using primers bearing a T7 promoter (Supplementary information, Table1).

***Reverse-transcription recombinase aided amplification (RT-RAA) assays***

Reverse-transcription recombinase aided amplification (RT-RAA) kit were purchased from

Hangzhou ZC Bio-Sci&Tech Co, Ltd and used according to the manufacturer's protocol with addition of 120 U Murine RNase Inhibitor (Vazyme). The 50 μL RT-RAA reaction system containing varying amounts of DNA input was incubated in 42°C for 30 minutes. All RAA products were directly used in the 55 μL detection assay as mentioned below.

***CDetection assays***

Reporter length optimization were performed with 30 nM AaCas12b, 36 nM sgRNA, 40 nM activator, 200 nM custom synthesized homopolymer ssDNA FQ reporter (Supplementary information, Table1) and NEBuffer™ 2.1 in a 20 μL reaction in a Corning® 384-well Polystyrene NBS Microplate. Reactions were incubated at 42°C for indicated time course in a fluorescence plate reader (BioTek Synergy 4) with fluorescent kinetics measured every 5 min ($\lambda_{ex}$=485 nm; $\lambda_{em}$=528 nm, transmission gain=61). The fluorescence results were analyzed by SigmaPlot software.

RdRp detection assays were performed as follow. 6 ul templates were RT-RAA amplified with CDetection system on lid. CDetection system consisted of 30 nM AaCas12b, 108 nM sgRNA (unless otherwise indicated), 200 nM custom synthesized homopolymer ssDNA FQ reporter (Supplementary information, Table1), 40 U Murine RNase Inhibitor (Vazyme), 10 mM Tris-HCl (pH 7.5), 10 mM $MgCl_2$ and 1 mM DTT in a 5 μL reaction. After finishing RT-RAA assay at 42°C for 30 minutes, CDetection was spun down and incubated at 42°C for 30 minutes in Applied Biosystems 7500 real-time PCR system (Thermo Fisher) with fluorescence kinetics measured every minute. ΔRn value were exported and analyzed by SigmaPlot software.

***Production of pseudovirus***

Blunt-SARS-CoV-2-RdRp, Blunt-SARS-CoV-RdRp, Blunt-MERS-CoV-RdRp plasmids were digested by BamHI (NEB) and XbaI (NEB), and ligated to lenti-CRISPR-V2 (a gift from Feng Zhang, Addgene plasmid #52961), to produce lenti-SARS-CoV-2-RdRp, lenti-SARS-CoV-RdRp and lenti-MERS-CoV-RdRp, respectively. Together with psPAX2 (a gift from Didier Trono, Addgene plasmid #12260) and pMD2.G (a gift from Didier Trono, Addgene plasmid #12259), lenti plasmids were co-transfected by lipofectamine LTX with Plus Reagent (Life Technology) into HEK293T cells. 48 and 72 hours after transfection, medium was harvested, filtered and concentrated by Amicon® Ultra-15 Centrifugal Filter (Millipore). Titration was calculated by QPCR.

### *Preparation of lentivirus*

To mimic practical virus detection, throat swab was added to virus transport medium (Youkang) and the resultant liquid was used to dilute lentivirus for our following experiments.

QIAamp Viral RNA Mini Kit (50) (QIAGEN) was used in accordance to the manufacturer's protocol. RNA extraction was done using 140 μL of sample and 25 μL nuclease-free water was used for elution.

Samples were treated with DTT/EDTA and heating prior to detection. 83 mM DTT and 0.83 mM EDTA were added to sample and followed by 2-step inactivation of 50℃ for 5 minutes and 64℃ for 5 minutes using a dry heat block. This product was used for subsequent CASdetec reaction.

### *Statistical analysis*

Statistical analyses were performed using Prism Software (GraphPad). For statistical comparison, Student's t test was employed. A value of $p < 0.05$ was considered significant.
