## Supplementary information, Fig. S for "SARS-CoV-2 detection with CRISPR diagnostics"

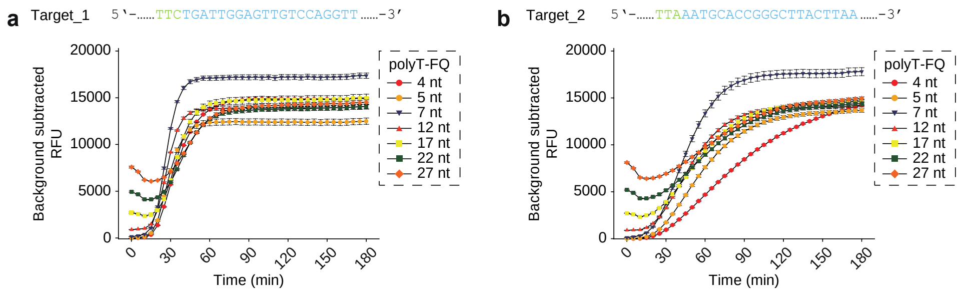


**Supplementary information, Fig. S1 Optimization of reporter length. a-b** Length preference of CDetection reporter. PAM sequences are colored in green, protospacers are colored in blue. Error bars indicate standard errors of the mean (s.e.m.)., n = 3. polyT-FQ, reporter made of T-homopolymer with FAM fluorophore and BHQ1 quencher.

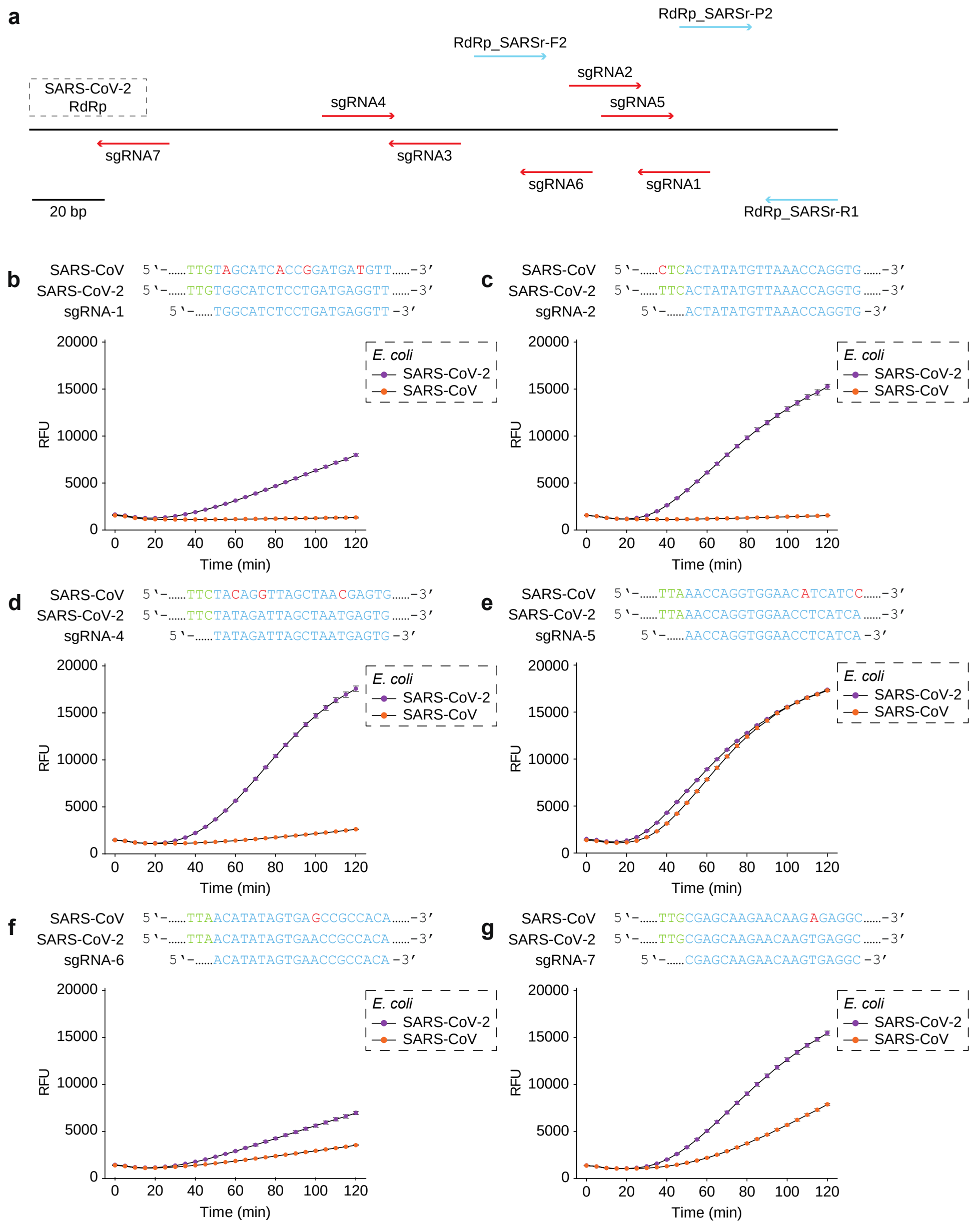


**Supplementary information, Fig. S2 SgRNA selection for SARS-CoV-2 detection. a** Schematics showing RdRp locus with sgRNAs, together with QPCR primers and probe disclosed by WHO. sgRNAs designed by ourselves are colored in red, QPCR primers and probe from WHO are colored in blue. bp, base pair. **b-g** Fluorescence kinetics of different sgRNAs for RdRp detection. *E. coli* cells bearing Blunt-SARS-CoV-RdRp or Blunt-SARS-CoV-2-RdRp were pre-incubated at 95°C for 10 min and used as templates for RAA and CDetection. PAM sequences are colored in green, protospacers are colored in blue, base pair mismatches are colored in red. Error bars indicate standard errors of the mean (s.e.m.), n = 3. RFU, relative fluorescence units.

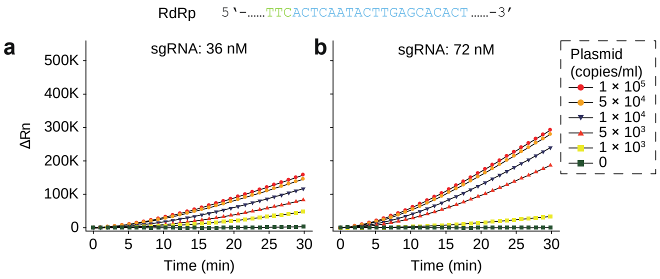


**Supplementary information, Fig. S3 Optimization of sgRNA concentration. a, b** Fluorescence kinetics of RdRp detection under 36 nM or 72 nM sgRNA-3. Plasmid bearing SARS-CoV-2-RdRp was serially diluted as shown in the legend. PAM sequences are colored in green, protospacers are colored in blue. $n$ = 2. ΔRn, ΔFluorescence, which refers to the Rn value of an experimental reaction minus the Rn value of the baseline signal generated by ABI 7500.

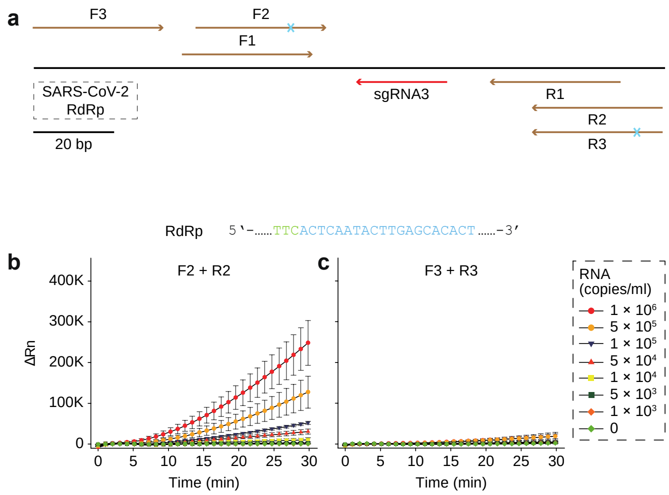


**Supplementary information, Fig. S4 Primer selection for SARS-CoV-2 detection. a** Schematics showing RdRp locus with RAA primers and sgRNA-3. The selected sgRNA-3 are colored in red, designed RAA primers are colored in brown. Blue cross indicated one base pair mismatch. bp, base pair. **b** Fluorescence kinetics of F2 and R2 based RdRp detection. SARS-CoV-2-RdRp RNA was serially diluted as shown in the legend. PAM sequences are colored in green, protospacers are colored in blue. Error bars indicate standard errors of the mean (s.e.m.), $n = 3$. ΔRn, ΔFluorescence, which refers to the Rn value of an experimental reaction minus the Rn value of the baseline signal generated by ABI 7500. **c** Fluorescence kinetics of F3 and R3 based RdRp detection. SARS-CoV-2-RdRp RNA was serially diluted as the legend show. Error bars indicate s.e.m., $n = 3$.

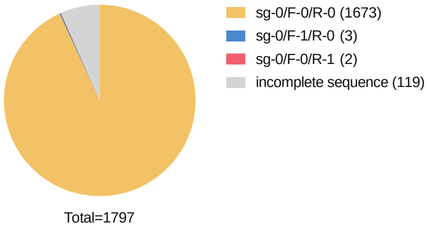


**Supplementary information, Fig. S5 Conservation analysis of selected sgRNA and primers.** Sequence alignment of 1797 reported SARS-CoV-2 sequences till March, 26$^{th}$, 2020. Sg-0/F-0/R-0, genomes which have no mismatches to sgRNA-3 or F1 or R1. Sg-0/F-1/R-0, genomes which have no mismatches to sgRNA-3 or R1, but 1 mismatch to F1. Sg-0/F-0/R-1, genomes which have no mismatches to sgRNA-3 or F1, but 1 mismatch to R1. Incomplete sequence, incomplete sequencing results which missed in the sgRNA-3, F1 and R1 region.

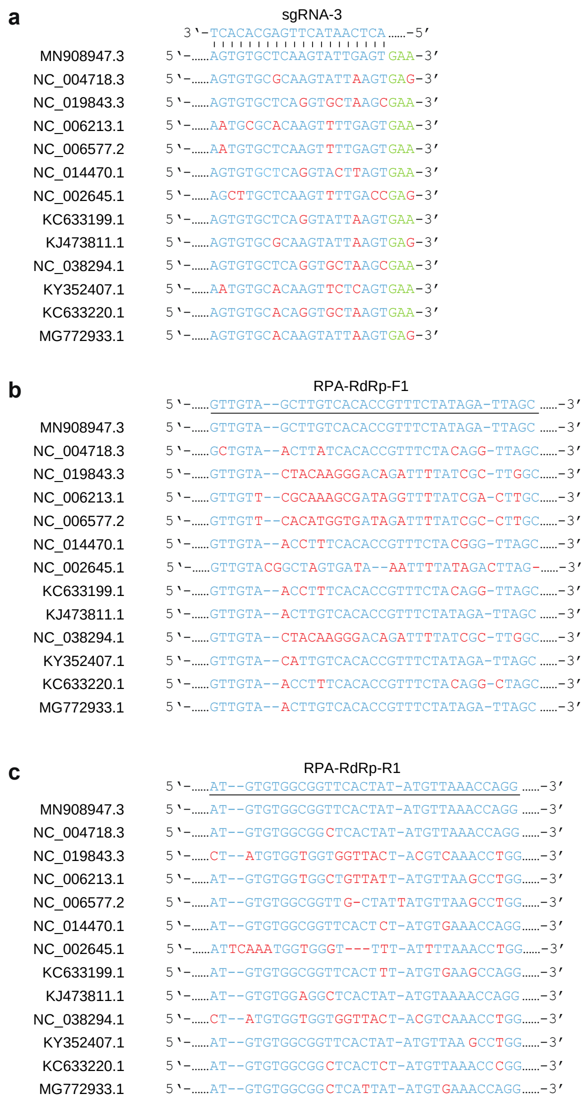


**Supplementary information, Fig. S6 Specificity analysis of selected sgRNA and primers.**

**a** Sequence alignment of selected sgRNA among typical coronavirus sequences. PAM sequences are colored in green, protospacers are colored in blue, mismatched bases are colored in red. **b** Sequence alignment of selected forward primer among typical coronavirus sequences. **c** Sequence alignment of selected reverse primer among typical coronavirus sequences.

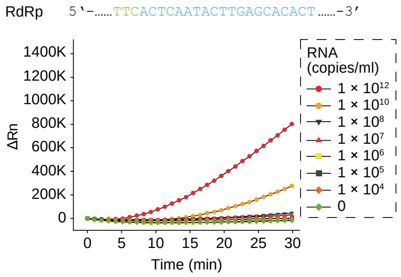


**Supplementary information, Fig. S7 One-step detection of SARS-CoV-2 fragments**. RAA and CDetection were combined into 1 step. SARS-CoV-2-RdRp RNA was serially diluted as the legend show. PAM sequences are colored in green, protospacers are colored in blue. $n = 2$. ΔRn, ΔFluorescence, which refers to the Rn value of an experimental reaction minus the Rn value of the baseline signal generated by ABI 7500.

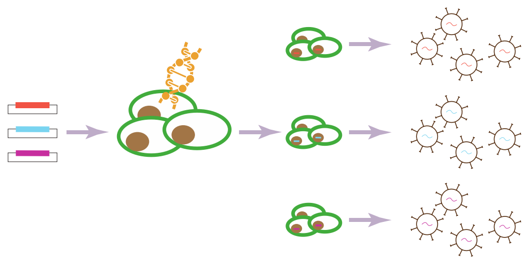

**Supplementary information, Fig. S8 Construction of pseudoviruses.** Schematics showing the procedure for pseudovirus packaging. Lentivirus plasmids bearing target RdRp region of SARS-CoV-2, SARS-CoV or MERS-CoV were co-transfected together with helper plasmids into HEK293T cells. Lentiviruses were harvested from the media.
