## Supplementary information, Table1 for "SARS-CoV-2 detection with CRISPR diagnostics"

**Supplementary information, Table 1 Nucleic acids used in this study.** The T7 promoter sequences are colored in green, and plasmid inserted sequences are colored in orange, respectively. Spacer sequences are underlined.

| | |
|---|---|
| **FQ ssDNA reporter** | |
| polyT-FQ-4nt | FAM-TTTT-BHQ1 |
| polyT-FQ-5nt | FAM-TTTTT-BHQ1 |
| polyT-FQ-7nt | FAM-TTTTTTT-BHQ1 |
| polyT-FQ-12nt | FAM-TTTTTTTTTTTT-BHQ1 |
| polyT-FQ-17nt | FAM-TTTTTTTTTTTTTTTTT-BHQ1 |
| polyT-FQ-22nt | FAM-TTTTTTTTTTTTTTTTTTTTTT-BHQ1 |
| polyT-FQ-27nt | FAM-TTTTTTTTTTTTTTTTTTTTTTTTTTT-BHQ1 |
| **Oligo** | |
| SARS-CoV-2-RdRp-1 | GTTTATAGTGATGTAGAAAACCCTCACCTTATGGGTTGGG |
| SARS-CoV-2-RdRp-2 | CATGGCTCTATCACATTTAGGATAATCCCAACCCATAAGGTGAGGGT |
| SARS-CoV-2-RdRp-3 | GGATTATCCTAAATGTGATAGAGCCATGCCTAACATGCTTAGAAT |
| SARS-CoV-2-RdRp-4 | GCAAGAACAAGTGAGGCCATAATTCTAAGCATGTTAGGCATGGCTCT |
| SARS-CoV-2-RdRp-5 | TTAGAATTATGGCCTCACTTGTTCTTGCTCGCAAACATACAACGT |
| SARS-CoV-2-RdRp-6 | GGTGTGACAAGCTACAACACGTTGTATGTTTGCGAGCAAGAACA |
| SARS-CoV-2-RdRp-7 | ACAACGTGTTGTAGCTTGTCACACCGTTTCTATAGATTAGCTAATGAG |
| SARS-CoV-2-RdRp-8 | CTCAATACTTGAGCACACTCATTAGCTAATCTATAGAAACGGTGTG |
| SARS-CoV-2-RdRp-9 | AGCTAATGAGTGTGCTCAAGTATTGAGTGAAATGGTCATGTGTGG |
| SARS-CoV-2-RdRp-10 | GTTTAACATATAGTGAACCGCCACACATGACCATTTCACTCAATAC |
| SARS-CoV-2-RdRp-11 | GTGTGGCGGTTCACTATATGTTAAACCAGGTGGAACCTCATCAGG |
| SARS-CoV-2-RdRp-12 | CATAAGCAGTTGTGGCATCTCCTGATGAGGTTCCACCTGGTTTAA |
| SARS-CoV-2-RdRp-13 | TCAGGAGATGCCACAACTGCTTATGCTAATAGTGTTTTTAACATTT |
| SARS-CoV-2-RdRp-14 | AAATGTTAAAAACACTATTAGCATAA |
| SARS-CoV-RdRp-1 | GTTTACAGTGATGTAGAAACTCCACACCTTATGGGTTGGGATTATCCAAAATGTG<br>AC |
| SARS-CoV-RdRp-2 | GCATGTTAGGCATGGCTCTGTCACATTTTGGATAATCCCAACCCATAAGG |
| SARS-CoV-RdRp-3 | GTGACAGAGCCATGCCTAACATGCTTAGGATAATGGCCTCTCTTGTTCTTGCTC |
| SARS-CoV-RdRp-4 | TGTGATAAGTTACAGCAAGTGTTATGTTTGCGAGCAAGAACAAGAGAGGCCATTA<br>TCCT |
| SARS-CoV-RdRp-5 | CGCAAACATAACACTTGCTGTAACTTATCACACCGTTTCTACAGGTTAGCTAACG |
| SARS-CoV-RdRp-6 | CCATCTCACTTAATACTTGCGCACACTCGTTAGCTAACCTGTAGAAACGGTGTGA |
| SARS-CoV-RdRp-7 | GAGTGTGCGCAAGTATTAAGTGAGATGGTCATGTGTGGCGGCTCACTATATG |
| SARS-CoV-RdRp-8 | CACCGGATGATGTTCCACCTGGTTTAACATATAGTGAGCCGCCACACATGAC |
| SARS-CoV-RdRp-9 | AAACCAGGTGGAACATCATCCGGTGATGCTACAACTGCTTATGCTAATAGTGTCT<br>TTAA |
| SARS-CoV-RdRp-10 | AAATGTTAAAGACACTATTAGCATAAGCAGTTGTAGC |
| MERS-CoV-RdRp-1 | TTGTACAAAGATGTTGATAATCCGCATCTTATGGGTTGGGATTACCCTAAGTGTG |
| MERS-CoV-RdRp-2 | GATTCTACACATATTAGGCATAGCTCTATCACACTTAGGGTAATCCCAACCCATA<br>AGA |
| MERS-CoV-RdRp-3 | GTGATAGAGCTATGCCTAATATGTGTAGAATCTTCGCTTCACTCATATTAGCTCG<br>TAA |
| MERS-CoV-RdRp-4 | CCTTGTAGTACAACAAGTGCCATGTTTACGAGCTAATATGAGTGAAGCGAAGATT<br>CT |

| | |
|---|---|
| MERS-CoV-RdRp-5 | CGTAAACATGGCACTTGTTGTACTACAAGGGACAGATTTTATCGCTTGGCAAATGAG |
| MERS-CoV-RdRp-6 | CGCTTAGCACCTGAGCACACTCATTTGCCAAGCGATAAAATCTGTCCCT |
| MERS-CoV-RdRp-7 | GAGTGTGCTCAGGTGCTAAGCGAATATGTTCTATGTGGTGGTGGTTACTACGTC |
| MERS-CoV-RdRp-8 | CCGCTACTGGTACCTCCAGGTTTGACGTAGTAACCACCACCACATAGAAC |
| MERS-CoV-RdRp-9 | AAACCTGGAGGTACCAGTAGCGGAGATGCCACCACTGCATATGCCAATAGTG |
| MERS-CoV-RdRp-10 | AAATGTTAAAGACACTATTGGCATATGCAGTGGTGGC |
| CoV-HKU1-1 | CTTATAAAGGATGTTGACAACCCTGTTCTTATGGGTTGGGATTATCCTAAATGTG |
| CoV-HKU1-2 | GCAAAATATTTGGCATAGCACGATCACATTTAGGATAATCCCAACCCATAAGA |
| CoV-HKU1-3 | GTGATCGTGCTATGCCAAATATTTTGCGTATTGTTAGTAGTTTAGTTTTGGCCC |
| CoV-HKU1-4 | CCATGTGAACAACAAAATTCATGTTTGCGGGCCAAAACTAAACTACTAACAATACG |
| CoV-HKU1-5 | CGCAAACATGAATTTTGTTGTTCACATGGTGATAGATTTTATCGCCTTGCGA |
| CoV-HKU1-6 | ATAACTATTTCACTCAAAACTTGAGCACATTCATTCGCAAGGCGATAAAATCTATCACC |
| CoV-HKU1-7 | GAATGTGCTCAAGTTTTGAGTGAAATAGTTATGTGTGGCGGTTGCTATTATG |
| CoV-HKU1-8 | CACTGCTAGTACCACCAGGCTTAACATAATAGCAACCGCCACACATAAC |
| CoV-HKU1-9 | AAGCCTGGTGGTACTAGCAGTGGTGATGCAACTACTGCTTTTGCTAATTCTGTTTTTAA |
| CoV-HKU1-10 | ATATATTAAAAACAGAATTAGCAAAAGCAGTAGTTGC |
| CoV-OC43-1 | GCCTTATTAAAGATGTTGACAATCCTGTACTTATGGGTTGGGATTATCCTAAG |
| CoV-OC43-2 | GGTTTGGCATAGCACGATCACACTTAGGATAATCCCAACCCATAAGTAC |
| CoV-OC43-3 | GTGTGATCGTGCTATGCCAAACCTACTACGTATTGTTAGTAGTTTGGTATTAGCCC |
| CoV-OC43-4 | CGAACAACATGTCTCATGTTTTCGGGCTAATACCAAACTACTAACAATACGTA |
| CoV-OC43-5 | CCCGAAAACATGAGACATGTTGTTCGCAAAGCGATAGGTTTTATCGACTTGCG |
| CoV-OC43-6 | CAATTTCACTCAAAACTTGTGCGCATTCATTCGCAAGTCGATAAAACCTATCGCTTT |
| CoV-OC43-7 | ATGAATGCGCACAAGTTTTGAGTGAAATTGTTATGTGTGGTGGCTGTTATTATGTTAAG |
| CoV-OC43-8 | CACTACTAGTGCCACCAGGCTTAACATAATAACAGCCACCACACATAACAA |
| CoV-OC43-9 | TTAAGCCTGGTGGCACTAGTAGTGGTGATGCAACTACTGCTTTTGCTAATTCAGTC |
| CoV-OC43-10 | ATGTTAAAGACTGAATTAGCAAAAGCAGTAGTTGCAT |
| CoV-NL63-1 | ATGAAACTTTTCTTGATTTTGCTTATTTTGCCCCTGGTTTCTTGCTTTTCT |
| CoV-NL63-2 | GTAACATAGAAATACTAGCATTACTGTTACATGTAGAAAAGCAAGAAACCAGGGGCAAA |
| CoV-NL63-3 | CATGTAACAGTAATGCTAGTATTTCTATGTTACAATTAGGTGTTCCTGATAACTCTT |
| CoV-NL63-4 | GCAACAAACCTGTGACAATAGTTGAAGAGTTATCAGGAACACCTAATTGTAACAT |
| CoV-NL63-5 | TCTTCAACTATTGTCACAGGTTTGTTGCCAGTCCATTGGATTTGTGCTAATC |
| CoV-NL63-6 | CATTGGCTGGGTAAGTAGATGTGCTCTGATTAGCACAAATCCAATGGACTGG |
| CoV-NL63-7 | AGCACATCTACTTACCCAGCCAATGGCTTTTTCTATATTGATGTCGGTAAACACCGTA |
| CoV-NL63-8 | CACTATGGAGTGCAAAAGCACTACGGTGTTTACCGACATCAATATAGAAAAA |

| | |
|---|---|
| CoV-NL63-9 | CGTAGTGCTTTTGCACTCCATAGTGGTTATTATGATGCTAACCAGTATTATATTTATCT |
| CoV-NL63-10 | TAGTGAGATAAATATAATACTGGTTAGCATCATAATA |
| CoV-229E-1 | GGTTCTCAAACAGTTCTAAGATGCGGTGATTGTTTACGCAGACCGATGTTGT |
| CoV-229E-2 | CATGATCATAGGCGCACTTAGTGCACAACATCGGTCTGCGTAAACAATCA |
| CoV-229E-3 | GCACTAAGTGCGCCTATGATCATGTGTTTGGCACTGATCATAAGTTCATTTTAGCTATT |
| CoV-229E-4 | GATGTGTTACACACATATGGTGTAATAGCTAAAATGAACTTATGATCAGTGCCAAA |
| CoV-229E-5 | GCTATTACACCATATGTGTGTAACACATCTGGCTGCAATGTAAATGACGTTAC |
| CoV-229E-6 | CAGTAATAATTCAAACCTCCAAGATACAGTTTTGTAACGTCATTTACATTGCAGCCAGA |
| CoV-229E-7 | AAACTGTATCTTGGAGGTTTGAATTATTACTGTGTAGACCACAAACCACATCT |
| CoV-229E-8 | CCAGCTGAACACAGTGGGAATGAAAGATGTGGTTTGTGGTCTACACAGTA |
| CoV-229E-9 | TCATTCCCACTGTGTTCAGCTGGTAATGTCTTTGGTTTGTACAAAAGTTCTGCTTTG |
| CoV-229E-10 | TGGAACCCAAAGCAGAACTTTTGTACAAACCAAAGAC |
| **DNA** | |
| Target_1_NTS-100 | AAACACTTACAGAAAGTTGTATTACCAGGTGGAAGGTTCTGATTGGAGTTGTCCAGGTTTTTGGCACGTTGAACAAATAATTGAACATCATGCATGAACA |
| Target_2_NTS-100 | CGCCAGGGTTTTCCCAGTCACGACAAAATCATAAAGTTAAATGCACCGGGCTTACTTAACAGCTTTTCGCTTTGAATCCTGTGTGAAATTGTTATCCGCT |
| ds_activator_RdRp_1 (277bp) | GTTTATAGTGATGTAGAAAACCCTCACCTTATGGGTTGGGATTATCCTAAATGTGATAGAGCCATGCCTAACATGCTTAGAATTATGGCCTCACTTGTTCTTGCTCGCAAACATACAACGTGTTGTAGCTTGTCACACCGTTTCTATAGATTAGCTAATGAGTGTGCTCAAGTATTGAGTGAAATGGTCATGTGTGGCGGTTCACTATATGTTAAACCAGGTGGAACCTCATCAGGAGATGCCACAACTGCTTATGCTAATAGTGTTTTTAACATTT |
| T7-sgRNA-Target_1 | TAATACGACTCACTATAGGGTCTAAAGGACAGAATTTTTCAACGGGTGTGCCAATGGCCACTTTCCAGGTGGCAAAGCCCGTTGAACTTCAAGCGAAGTGGCACTGATTGGAGTTGTCCAGGTT |
| T7-sgRNA-Target_2 | TAATACGACTCACTATAGGGTCTAAAGGACAGAATTTTTCAACGGGTGTGCCAATGGCCACTTTCCAGGTGGCAAAGCCCGTTGAACTTCAAGCGAAGTGGCACAATGCACCGGGCTTACTTAA |
| T7-AasgRNA-RdRp-1 | TAATACGACTCACTATAGGGTCTAAAGGACAGAATTTTTCAACGGGTGTGCCAATGGCCACTTTCCAGGTGGCAAAGCCCGTTGAACTTCAAGCGAAGTGGCACTGGCATCTCCTGATGAGGTT |
| T7-AasgRNA-RdRp-2 | TAATACGACTCACTATAGGGTCTAAAGGACAGAATTTTTCAACGGGTGTGCCAATGGCCACTTTCCAGGTGGCAAAGCCCGTTGAACTTCAAGCGAAGTGGCACACTATATGTTAAACCAGGTG |
| T7-AasgRNA-RdRp-3 | TAATACGACTCACTATAGGGTCTAAAGGACAGAATTTTTCAACGGGTGTGCCAATGGCCACTTTCCAGGTGGCAAAGCCCGTTGAACTTCAAGCGAAGTGGCACACTCAATACTTGAGCACACT |
| T7-AasgRNA-RdRp-4 | TAATACGACTCACTATAGGGTCTAAAGGACAGAATTTTTCAACGGGTGTGCCAATGGCCACTTTCCAGGTGGCAAAGCCCGTTGAACTTCAAGCGAAGTGGCACTATAGATTAGCTAATGAGTG |

| | |
|---|---|
| T7-AasgRNA-RdRp-5 | TAATACGACTCACTATAGGGTCTAAAGGACAGAATTTTTCAACGGGTGTGCCAATGGCCACTTTCCAGGTGGCAAAGCCCGTTGAACTTCAAGCGAAGTGGCACAACCAGGTGGAACCTCATCA |
| T7-AasgRNA-RdRp-6 | TAATACGACTCACTATAGGGTCTAAAGGACAGAATTTTTCAACGGGTGTGCCAATGGCCACTTTCCAGGTGGCAAAGCCCGTTGAACTTCAAGCGAAGTGGCACACATATAGTGAACCGCCACA |
| T7-AasgRNA-RdRp-7 | TAATACGACTCACTATAGGGTCTAAAGGACAGAATTTTTCAACGGGTGTGCCAATGGCCACTTTCCAGGTGGCAAAGCCCGTTGAACTTCAAGCGAAGTGGCACCGAGCAAGAACAAGTGAGGC |
| **Primers** | |
| Target_1-F | AAACACTTACAGAAAGTTGTATTACCAGGT |
| Target_1-R | TGTTCATGCATGATGTTCAATTATTTGTTC |
| Target_2-F | CGCCAGGGTTTTCCCAGTCACGAC |
| Target_2-R | AGCGGATAACAATTTCACACAGGA |
| RPA-RdRp-F0 | GTTTATAGTGATGTAGAAAACCCTCACCTTAT (used in Fig. 1a, Fig. S2b-h) |
| RPA-RdRp-R0 | AAATGTTAAAAACACTATTAGCATAAGCAGTT (used in Fig. 1a, Fig. S2b-h) |
| RPA-RdRp-F1 | GTTGTAGCTTGTCACACCGTTTCTATAGATTAGC |
| RPA-RdRp-R1 | CCTGGTTTAACATATAGTGAACCGCCACACAT |
| RPA-RdRp-F2 | TTGTAGCTTGTCACACCGTTTATATAGATTAG |
| RPA-RdRp-R2 | CACCTGGTTTAACATATAGTGAACCGCCACACA |
| RPA-RdRp-F3 | TTAGAATTATGGCCTCACTTGTTCTTGCTC |
| RPA-RdRp-R3 | CACCTGGATTAACATATAGTGAACCGCCACACA |
| T7-AasgRNA-F | TAATACGACTCACTATAGGGTCTAAAGGACAGAATTTTTCAACGGGTG |
| T7-sgRNA-Target_1-R | AACCTGGACAACTCCAATCAGTGCCACTTCGCTTGAAGTTCA |
| T7-sgRNA-Target_2-R | TTAAGTAAGCCCGGTGCATTGTGCCACTTCGCTTGAAGTTCA |
| T7-sgRNA-RdRp-1-R | AACCTCATCAGGAGATGCCAGTGCCACTTCGCTTGAAGTTCA |
| T7-sgRNA-RdRp-2-R | CACCTGGTTTAACATATAGTGTGCCACTTCGCTTGAAGTTCA |
| T7-sgRNA-RdRp-3-R | AGTGTGCTCAAGTATTGAGTGTGCCACTTCGCTTGAAGTTCA |
| T7-sgRNA-RdRp-4-R | CACTCATTAGCTAATCTATAGTGCCACTTCGCTTGAAGTTCA |
| T7-sgRNA-RdRp-5-R | TGATGAGGTTCCACCTGGTTGTGCCACTTCGCTTGAAGTTCA |
| T7-sgRNA-RdRp-6-R | TGTGGCGGTTCACTATATGTGTGCCACTTCGCTTGAAGTTCA |
| T7-sgRNA-RdRp-7-R | GCCTCACTTGTTCTTGCTCGGTGCCACTTCGCTTGAAGTTCA |
| QPCR-lenti-F | GGACGTCCTTCTGCTACGTC(used for determination of pseudovirus titration) |
| QPCR-lenti-R | GAGATCCGACTCGTCTGAGG(used for determination of pseudovirus titration) |
| **AasgRNA** | |
| AasgRNA-Target_1 | GUCUAAAGGACAGAAUUUUUCAACGGGUGUGCCAAUGGCCACUUUCCAGGUGGCAAAGCCCGUUGAACUUCAAGCGAAGUGGCACUGAUUGGAGUUGUCCAGGUU |
| AasgRNA-Target_2 | GUCUAAAGGACAGAAUUUUUCAACGGGUGUGCCAAUGGCCACUUUCCAGGUGGCAAAGCCCGUUGAACUUCAAGCGAAGUGGCACAAUGCACCGGGCUUACUUAA |
| AasgRNA-RdRp-1 | GUCUAAAGGACAGAAUUUUUCAACGGGUGUGCCAAUGGCCACUUUCCAGGUGGCAAAGCCCGUUGAACUUCAAGCGAAGUGGCACUGGCAUCUCCUGAUGAGGUU |
| AasgRNA-RdRp-2 | GUCUAAAGGACAGAAUUUUUCAACGGGUGUGCCAAUGGCCACUUUCCAGGUGGCAAAGCCCGUUGAACUUCAAGCGAAGUGGCACUAUAUGUUAAACCAGGUG |
| AasgRNA-RdRp-3 | GUCUAAAGGACAGAAUUUUUCAACGGGUGUGCCAAUGGCCACUUUCCAGGUGGCAAAGCCCGUUGAACUUCAAGCGAAGUGGCACACUCAAUACUUGAGCACACU |
| AasgRNA-RdRp-4 | GUCUAAAGGACAGAAUUUUUCAACGGGUGUGCCAAUGGCCACUUUCCAGGUGGCAAAGCCCGUUGAACUUCAAGCGAAGUGGCACUAUAGAUUAGCUAAUGAGUG |

| | |
|---|---|
| AasgRNA-RdRp-5 | GUCUAAAGGACAGAAUUUUUCAACGGGUGUGCCAAUGGCCACUUUCCAGGUGGCAAAGCCCGUUGAACUUCAAGCGAAGUGGCACAACCAGGUGGAACCUCAUCA |
| AasgRNA-RdRp-6 | GUCUAAAGGACAGAAUUUUUCAACGGGUGUGCCAAUGGCCACUUUCCAGGUGGCAAAGCCCGUUGAACUUCAAGCGAAGUGGCACACAUAUAGUGAACCGCCACA |
| AasgRNA-RdRp-7 | GUCUAAAGGACAGAAUUUUUCAACGGGUGUGCCAAUGGCCACUUUCCAGGUGGCAAAGCCCGUUGAACUUCAAGCGAAGUGGCACCGAGCAAGAACAAGUGAGGC |
| **pEASY-Blunt plasmid** | |
| pEASY-Blunt-SARS-CoV-2-RdRp | ...ATATCTGCAGAATTGCCCTTGTTTATAGTGATGTAGAAAACCCTCACCTTATGGGTTGGGATTATCCTAAATGTGATAGAGCCATGCCTAACATGCTTAGAATTATGGCCTCACTTGTTCTTGCTCGCAAACATACAACGTGTTGTAGCTTGTCACACCGTTTCTATAGATTAGCTAATGAGTGTGCTCAAGTATTGAGTGAAATGGTCATGTGTGGCGGTTCACTATATGTTAAACCAGGTGGAACCTCATCAGGAGATGCCACAACTGCTTATGCTAATAGTGTTTTTAACATTTAAGGGCAATTCCAGCACACT |
| pEASY-Blunt-SARS-CoV-RdRp | ...ATATCTGCAGAATTGCCCTTGTTTACAGTGATGTAGAAACTCCACACCTTATGGGTTGGGATTATCCAAAATGTGACAGAGCCATGCCTAACATGCTTAGGATAATGGCCTCTCTTGTTCTTGCTCGCAAACATAACACTTGCTGTAACTTATCACACCGTTTCTACAGGTTAGCTAACGAGTGTGCGCAAGTATTAAGTGAGATGGTCATGTGTGGCGGCTCACTATATGTTAAACCAGGTGGAACATCATCCGGTGATGCTACAACTGCTTATGCTAATAGTGTCTTTAACATTTAAGGGCAATTCCAGCACACT |
| pEASY-Blunt-MERS-CoV-RdRp | ...ATATCTGCAGAATTGCCCTTAAATGTTAAAGACACTATTGGCATATGCAGTGGTGGCATCTCCGCTACTGGTACCTCCAGGTTTGACGTAGTAACCACCACCACATAGAACATATTCGCTTAGCACCTGAGCACACTCATTTGCCAAGCGATAAAATCTGTCCCTTGTAGTACAACAAGTGCCATGTTTACGAGCTAATATGAGTGAAGCGAAGATTCTACACATATTAGGCATAGCTCTATCACACTTAGGGTAATCCCAACCCATAAGATGCGGATTATCAACATCTTTGTACAAATTCCAGCACACTGGCGGCC... |
| pEASY-Blunt-CoV-HKU1-RdRp | ...ATATCTGCAGAATTGCCCTTATATATTAAAAACAGAATTAGCAAAAGCAGTAGTTGCATCACCACTGCTAGTACCACCAGGCTTAACATAATAGCAACCGCCACACATAACTATTTCACTCAAAACTTGAGCACATTCATTCGCAAGGCGATAAAATCTATCACCATGTGAACAACAAAATTCATGTTTGCGGGCCAAAACTAAACTACTAACAATACGCAAAATATTTGGCATAGCACGATCACATTTAGGATAATTCCAGCACACTGGCGGCCGT... |
| pEASY-Blunt-CoV-OC43-RdRp | ...ATATCTGCAGAATTGCCCTTATGTTAAAGACTGAATTAGCAAAAGCAGTAGTTGCATCACCACTACTAGTGCCACCAGGCTTAACATAATAACAGCCACCACACATAACAATTTCACTCAAAACTTGTGCGCATTCATTCGCAAGTCGATAAAACCTATCGCTTTGCGAACAACATGTCTCATGTTTTCGGGCTAATACCAAACTACTAACAATACGTAGTAGGTTTGGCATAGCACGATCACACTTAGGATAATCCCAACCCATAAGTACAGGATTGTCAACATCTTTAATAAGGCAAGGGCAATTCCAGCACACT... |
| pEASY-Blunt-CoV-NL63-RdRp | ...ATATCTGCAGAATTGCCCTTTAGTGAGATAAATATAATACTGGTTAGCATCATAATAACCACTATGGAGTGCAAAAGCACTACGGTGTTTACCGACATCAATATAGAAAAAGCCATTGGCTGGGTAAGTAGATGTGCTCTGATTAGCACAAATCCAATGGACTGGCAACAAACCTGTGACAATAGTTGAAGAGTTATCAGGAACACCTAATTGTAACATAGAAATACTAGCATTACTGTTACATGTAGAAAAGCAAGAAACCAGGGGCAAAATAAGCAAAATCAAGAAAAGTTTCATAAGGGCAATTCCAGCACACT... |
| pEASY-Blunt-CoV-229E-RdRp | ...ATATCTGCAGAATTGCCCTTTGGAACCCAAAGCAGAACTTTTGTACAAACCAAAGACATTACCAGCTGAACACAGTGGGAATGAAAGATGTGGTTTGTGGTCTACACAGTAATAATTCAAACCTCCAAGATACAGTTTTGTAACGTCATTTACATTGCAGCCAGATGTGTTACACACATATGGTGTAATAGCTAAAATGAACTTATGATCAGTGCCA |

| | |
|---|---|
| | AACACATGATCATAGGCGCACTTAGTGCACAACATCGGTCTGCGTAAACAATCAC<br>CGCATCTTAGAACTGTTTGAGAACCAAGGGCAATTCCAGCACACT... |
| **RNA** | |
| SARS-CoV-2-RdRp_RNA | GUUUAUAGUGAUGUAGAAAACCCUCACCUUAUGGGUUGGGAUUAUCCUAAAUGUG<br>AUAGAGCCAUGCCUAACAUGCUUAGAAUUAUGGCCUCACUUGUUCUUGCUCGCAA<br>ACAUACAACGUGUUGUAGCUUGUCACACCGUUUCUAUAGAUUAGCUAAUGAGUGU<br>GCUCAAGUAUUGAGUGAAAUGGUCAUGUGUGGCGGUUCACUAUAUGUUAAACCAG<br>GUGGAACCUCAUCAGGAGAUGCCACAACUGCUUAUGCUAAUAGUGUUUUUAACAU<br>UU |
| SARS-CoV-RdRp_RNA | GUUUACAGUGAUGUAGAAACUCCACACCUUAUGGGUUGGGAUUAUCCAAAAUGUG<br>ACAGAGCCAUGCCUAACAUGCUUAGGAUAAUGGCCUCUCUUGUUCUUGCUCGCAA<br>ACAUAACACUUGCUGUAACUUAUCACACCGUUUCUACAGGUUAGCUAACGAGUGU<br>GCGCAAGUAUUAAGUGAGAUGGUCAUGUGUGGCGGCUCACUAUAUGUUAAACCAG<br>GUGGAACAUCAUCCGGUGAUGCUACAACUGCUUAUGCUAAUAGUGUCUUUAACAU<br>UU |
| MERS-CoV-RdRp_RNA | AAAUGUUAAAGACACUAUUGGCAUAUGCAGUGGUGGCAUCUCCGCUACUGGUACC<br>UCCAGGUUUGACGUAGUAACCACCACCACAUAGAACAUAUUCGCUUAGCACCUGA<br>GCACACUCAUUUGCCAAGCGAUAAAAUCUGUCCCUUGUAGUACAACAAGUGCCAU<br>GUUUACGAGCUAAUAUGAGUGAAGCGAAGAUUCUACACAUAUUAGGCAUAGCUCU<br>AUCACACUUAGGGUAAUCCCAACCCAUAAGAUGCGGAUUAUCAACAUCUUUGUAC<br>AA |
| CoV-HKU1-RdRp_RNA | AUAUAUUAAAAACAGAAUUAGCAAAAGCAGUAGUUGCAUCACCACUGCUAGUACC<br>ACCAGGCUUAACAUAAUAGCAACCGCCACACAUAACUAUUUCACUCAAAACUUGA<br>GCACAUUCAUUCGCAAGGCGAUAAAAUCUAUCACCAUGUGAACAACAAAAUUCAU<br>GUUUGCGGGCCAAAACUAAACUACUAACAAUACGCAAAAUAUUUGGCAUAGCACG<br>AUCACAUUUAGGAUAAU |
| CoV-OC43-RdRp_RNA | AUGUUAAAGACUGAAUUAGCAAAAGCAGUAGUUGCAUCACCACUACUAGUGCCAC<br>CAGGCUUAACAUAAUAACAGCCACCACACAUAACAAUUUCACUCAAAACUUGUGC<br>GCAUUCAUUCGCAAGUCGAUAAAACCUAUCGCUUUGCGAACAACAUGUCUCAUGU<br>UUUCGGGCUAAUACCAAACUACUAACAAUACGUAGUAGGUUUGGCAUAGCACGAU<br>CACACUUAGGAUAAUCCCAACCCAUAAGUACAGGAUUGUCAACAUCUUUAAUAAG<br>GC |
| CoV-NL63-RdRp_RNA | UAGUGAGAUAAAUAUAAUACUGGUUAGCAUCAUAAUAACCACUAUGGAGUGCAAA<br>AGCACUACGGUGUUUACCGACAUCAAUAUAGAAAAAGCCAUUGGCUGGGUAAGUA<br>GAUGUGCUCUGAUUAGCACAAAUCCAAUGGACUGGCAACAAACCUGUGACAAUAG<br>UUGAAGAGUUAUCAGGAACACCUAAUUGUAACAUAGAAAUACUAGCAUUACUGUU<br>ACAUGUAGAAAAGCAAGAAACCAGGGGCAAAAUAAGCAAAAUCAAGAAAAGUUUC<br>AU |
| CoV-229E-RdRp_RNA | UGGAACCCAAAGCAGAACUUUUGUACAAACCAAAGACAUUACCAGCUGAACACAG<br>UGGGAAUGAAAGAUGUGGUUUGUGGUCUACACAGUAAUAAUUCAAACCUCCAAGA<br>UACAGUUUUGUAACGUCAUUUACAUUGCAGCCAGAUGUGUUACACACAUAUGGUG<br>UAAUAGCUAAAAUGAACUUAUGAUCAGUGCCAAACACAUGAUCAUAGGCGCACUU<br>AGUGCACAACAUCGGUCUGCGUAAACAAUCACCGCAUCUUAGAACUGUUUGAGAA<br>CC |
| **Lentivirus plasmid** | |
| Lenti-SARS-CoV-2-RdRp | ...AGTGTGCTGGAATTGCCCTTGTTTATAGTGATGTAGAAAACCCTCACCTTAT<br>GGGTTGGGATTATCCTAAATGTGATAGAGCCATGCCTAACATGCTTAGAATTATG<br>GCCTCACTTGTTCTTGCTCGCAAACATACAACGTGTTGTAGCTTGTCACACCGTT |

| | |
|---|---|
| | TCTATAGATTAGCTAATGAGTGTGCTCAAGTATTGAGTGAAATGGTCATGTGTGG<br>CGGTTCACTATATGTTAAACCAGGTGGAACCTCATCAGGAGATGCCACAACTGCT<br>TATGCTAATAGTGTTTTTAACATTTAAGGGCAATTCTGCAGATAT... |
| Lenti-SARS-CoV-RdRp | ...ATATCTGCAGAATTGCCCTTGtttacagtgatgtagaaactccacaccttat<br>gggttgggattatccaaaatgtgacagagccatgcctaacatgcttaggataatg<br>gcctctcttgttcttgctcgcaaacataacacttgctgtaacttatcacaccgtt<br>tctacaggttagctaacgagtgtgcgcaagtattaagtgagatggtcatgtgtgg<br>cggctcactatatgttaaaccaggtggaacatcatccggtgatgctacaactgct<br>tatgctaatagtgtctttaacatttAAGGGCAATTCCAGCACACT... |
| Lenti-MERS-CoV-RdRp | ...AACACAGGACCGGTTCTAGAttgtacaaagatgttgataatccgcatcttat<br>gggttgggattaccctaagtgtgatagagctatgcctaatatgtgtagaatcttc<br>gcttcactcatattagctcgtaaacatggcacttgttgtactacaagggacagat<br>tttatcgcttggcaaatgagtgtgctcaggtgctaagcgaatatgttctatgtgg<br>tggtggttactacgtcaaacctggaggtaccagtagcggagatgccaccactgca<br>tatgccaatagtgtctttaacatttAAGGGCAATTCTGCAGATAT... |
